# Data quality biases normative models derived from fetal brain MRI

**DOI:** 10.64898/2026.01.22.700996

**Authors:** Thomas Sanchez, Angeline Mihailov, Gerard Martí-Juan, Nadine Girard, Aurélie Manchon, Mathieu Milh, Elisenda Eixarch, Vincent Dunet, Mériam Koob, Léo Pomar, Joanna Sichitiu, Miguel A. González Ballester, Oscar Camara, Gemma Piella, Meritxell Bach Cuadra, Guillaume Auzias

## Abstract

Normative modeling is increasingly used to characterize typical growth trajectories and identify atypical neurodevelopment, including early brain development using magnetic resonance imaging (MRI) acquired before birth. Recent work has emphasized the importance of large sample sizes for accurate and robust centile estimation. In this study, we investigate how image quality influences fetal brain normative models, a critical factor in this context where MRI is acquired on a moving fetus *in utero*.

Using a multi-centric cohort of 635 fetal MRI scans, we applied a standardized visual quality control (QC) protocol with continuous quality ratings. We fit normative models for multiple brain structures under progressively relaxed QC stringency, and quantified the deviations in centile estimates relative to a high-quality reference subgroup. Our results showed that including lower-quality data systematically biased normative centiles, with the strongest effects observed in the outer centiles, particularly the lower tail (1st–10th). Bias increased progressively as QC stringency was relaxed and could not be attributed solely to the number of subjects used to fit the models. Quality-induced bias was structure-dependent, and often not visually apparent at the segmentation level.

These findings highlight that image quality is an important source of bias in normative fetal brain modeling, and that increasing sample size at the expense of quality may systematically affect centile estimates, potentially jeopardizing the utility of the model.

## 1 Introduction

Accurate and robust estimation of normative trajectories of brain development is crucial for detecting atypical neurodevelopmental processes and potential neurological disorders (Bethlehem et al., 2022; Fraza et al., 2025; Kyriakopoulou et al., 2017; Rutherford et al., 2022). Structural magnetic resonance imaging (MRI) has become increasingly used to study the developing brain, offering a non-invasive means to characterize typical and atypical neurodevelopmental trajectories throughout gestation and into early postnatal life (Bethlehem et al., 2022; Ji et al., 2025; Kyriakopoulou et al., 2017; Machado-Rivas et al., 2022; Mihailov et al., 2025). However, the reliability and validity of measures derived from fetal MRI directly depend on the quality of the acquired images, which can be affected by fetal motion, maternal physiological noise, and scanner-related artifacts. Rigorous QC is particularly critical in fetal imaging, where rapid anatomical changes throughout gestational age interact with complex reconstruction and segmentation pipelines, potentially introducing subtle and structure-specific artifacts (Sanchez et al., 2024).

While quality control (QC) protocols are now well established in adult brain MRI (Alfaro-Almagro et al., 2018; Backhausen et al., 2016; Esteban et al., 2017; Kim et al., 2025; Monereo-Sánchez et al., 2021), this is not the case in fetal brain imaging. In this field, qualitative visual inspection of each scan remains common practice to exclude images with severe artifacts or poor contrast (Ducharme et al., 2016; Namburete et al., 2023; Rosen et al., 2018). Despite the major potential impact of QC on the reported results, the description of the operational criteria are often insufficient to enable other groups to reproduce the full procedure, which impedes both the comparison and reproducibility of results across studies. In addition, while most studies report excluding low-quality data, the impact of varying degrees of data quality—beyond binary inclusion or exclusion—on normative modeling results is rarely investigated. More specifically, the impact of sub-optimal data quality on the estimation of fetal brain normative trajectories and centiles has not been investigated. In this work, we address the following questions:

1. How does the inclusion of sub-optimal quality MRI scans bias the estimation of median and outer centiles in fetal brain normative models?
2. To what extent is this bias driven by image quality itself, rather than to changes in effective sample size induced by quality-based data exclusion?

To answer these questions, we investigate the effect of varying visual QC stringency on normative centile predictions for multiple fetal brain structures, using a large multi-centric cohort of 635 fetal MRI scans. Leveraging a recently released standardized visual QC protocol (Bach Cuadra et al., 2025; Sanchez et al., 2024) that provides continuous image-level quality ratings, we define nested QC subgroups of increasing stringency and quantify the resulting centile bias relative to a high-quality reference subgroup. We further disentangle the respective contributions of data quality and sample size by means of a weighted bootstrapping strategy that allows independent control over sample size and average data quality. Together, these analyses aim to characterize how data quality systematically influences fetal brain normative models, beyond simple inclusion–exclusion decisions, and to assess the extent to which increasing sample size can compensate for lower data quality.

## 2 Methods

### 2.1 Data

#### 2.1.1 Cohorts and demographics

Brain MRI examinations were retrospectively collected from ongoing clinical research studies in three hospitals: La Timone, Marseille, France (n=216); BCNatal (Hospital Clínic and Hospital Sant Joan de Déu), Barcelona, Spain (n=91); Lausanne University Hospital (CHUV), Switzerland (n=36), along with a research dataset, the fetal dHCP (n=292). Exclusion criteria included twin pregnancies and any pathology or malformation in the fetal MRI scans. Ethical approval was obtained from the institutional review boards of all participating centers (La Timone: Aix-Marseille University N°2022-04-14-003; BCNatal: HCB/2022/0533; CHUV: CER-VD 2021-00124).

##### Clinical dataset

The clinical fetal MRI data were acquired with different Siemens Healthineers scanners (Erlangen, Germany) at 1.5 tesla (T) or 3T across hospitals. The fetal brain MRI protocol included T2w HASTE (Half-Fourier Acquisition Single-shot Turbo spin Echo imaging) sequences acquired in at least three orthogonal directions (axial, coronal, sagittal). Table 1 highlights the demographic information of the dataset.

**Table 1:**
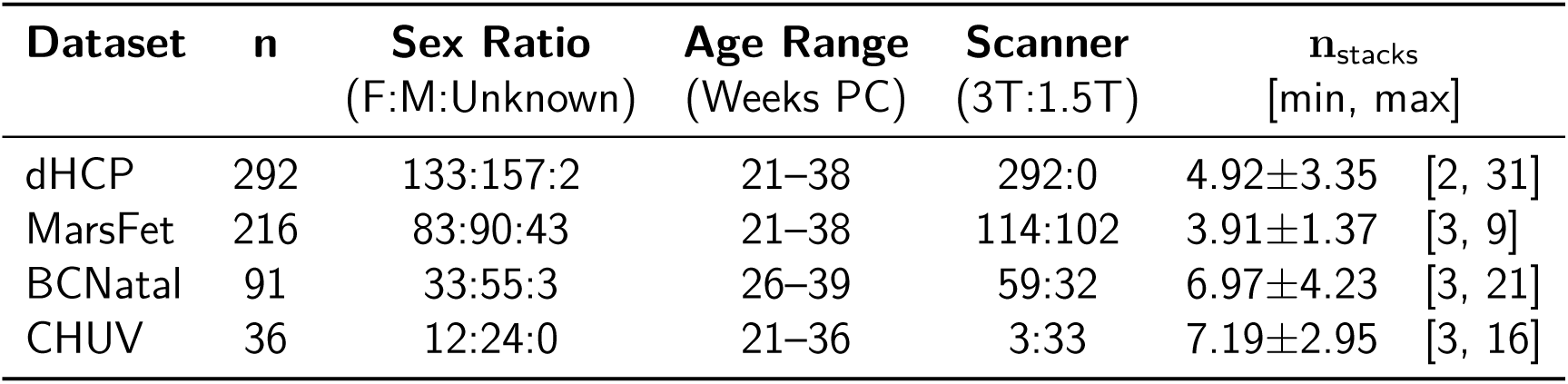
Summary of fetal MRI datasets.

| Dataset | n | Sex Ratio<br>(F:M:Unknown) | Age Range<br>(Weeks PC) | Scanner<br>(3T:1.5T) | $n_{\text{stacks}}$<br>[min, max] |
| --- | --- | --- | --- | --- | --- |
| dHCP | 292 | 133:157:2 | 21–38 | 292:0 | $4.92 \pm 3.35$ [2, 31] |
| MarsFet | 216 | 83:90:43 | 21–38 | 114:102 | $3.91 \pm 1.37$ [3, 9] |
| BCNatal | 91 | 33:55:3 | 26–39 | 59:32 | $6.97 \pm 4.23$ [3, 21] |
| CHUV | 36 | 12:24:0 | 21–36 | 3:33 | $7.19 \pm 2.95$ [3, 16] |

To establish a healthy fetal cohort, a multidisciplinary team, including specialists in neuroradiology, obstetrics, and pediatric neurology, formalized a list of exclusion criteria to define standardized clinical and anatomical normality across sites. The detailed inclusion and exclusion criteria are discussed in the Supplementary Table S1. Applying these criteria led to a cohort of 635 subjects (261:326:46 (F:M:unknown), age range between 20.85 - 38.86 weeks of gestational age). We now describe these datasets in detail.

1. MarsFet. The MarsFet fetal dataset was assembled retrospectively from MRI examinations performed during routine prenatal clinical care at La Timone Hospital in Marseille between 2008 and 2021. The data were acquired on two Siemens scanners (Skyra 3T and SymphonyTim 1.5T) with at least three stacks acquired in orthogonal orientations. Spatial resolution was between 0.6 − 0.7 mm in-plane and slice thickness between 3 − 3.6 mm. After applying predefined exclusion criteria, 216 fetal participants were retained for analysis.
2. BCNatal. The BCNatal dataset was retrospectively assembled from fetal MRI scans of uncomplicated pregnancies acquired at Hospital Clínic Barcelona and Hospital Sant Joan de Déu, Spain, grouped around 28 and 34 weeks of gestational age (GA), where MRI examinations are typically performed. The data were acquired on two Siemens scanners (TrioTim 3T and Aera 1.5T), with a minimum of four stacks per subject, including at least one in each orthogonal plane. Spatial resolution was 0.51 × 0.51 × 3.5 mm^3^ at 3T and 0.55 × 0.55 × 3.6 mm^3^ at 1.5T. After applying predefined exclusion criteria, 91 fetal participants were included in the analysis.
3. CHUV. The CHUV dataset was assembled retrospectively from MRI examinations performed during standard prenatal clinical care between 2013 and 2024. The data were acquired on four Siemens scanners (Skyra and Skyra_fit at 3T, and Aera and MAGNETOM Sola at 1.5T), with a minimum of six stacks (at least two for each orthogonal plane). Spatial resolution was 0.55 × 0.55 × 3 mm^3^ at 3T, and 1.1 × 1.1 × 3.3 mm^3^ at 1.5T. After applying predefined exclusion criteria, 36 fetal participants were retained for analysis.

##### Research dataset

###### Fetal dHCP

The fetal dHCP dataset was obtained from the fourth release of the publicly available developing Human Connectome Project (dHCP)^1^. This dataset consists of 297 MRI sessions from 273 fetuses. As detailed in (Price et al., 2019), MRI data were acquired on a 3T Philips Achieva scanner. Structural T2w data was acquired from 6 uniquely oriented stacks centered to the fetal brain using a zoomed multiband single-shot TSE sequence, at an in-plane resolution of 1.1 mm and slice thickness of 2.2 mm. This dataset resulted in a final sample of 292 fetal participants.

### 2.2 Data processing

To ensure reproducible processing across the datasets, we relied on Fetpype, an open source image processing pipeline designed to standardize the pre-processing, reconstruction and segmentation of fetal brain MRI (Sanchez et al., 2025)^2^.

#### Super-resolution reconstruction

Clinical fetal brain MRI acquisition are typically acquired using multiple orientations with a good in-plane resolution and thick slices. We reconstructed these data into a single high-resolution volume using state-of-the art super-resolution reconstruction (SRR) methods integrated in Fetpype. The preprocessing and reconstruction in Fetpype consists of the following steps: brain extraction using Fetal-BET (Faghihpirayesh et al., 2024), non-local means denoising (Manjón et al., 2008) and N4 bias-field correction Tustison et al., 2010, 3D volume reconstruction using NeSVoR (v.0.5.0) J. Xu et al., 2023, including all available HASTE stacks to reconstruct each subject (no selection of good quality stacks). Data were reconstructed at 0.5mm isotropic for MarsFet and BCNatal, and 0.8mm isotropic at CHUV.

For the fetal dHCP, we used the released 3D isotropic (0.5 mm iso) MRI scans that were reconstructed using the method described in (Kuklisova-Murgasova et al., 2012; A. U. Uus et al., 2022).

#### Segmentation

All data were then segmented using BOUNTI (A. U. Uus et al., 2023), which is integrated in Fetpype. BOUNTI’s segmentation features 19 different brain regions. For both hemispheres, extra-cerebral CSF (eCSF), cortical gray matter (cGM), white matter (WM), lateral ventricles (LV), cerebellum, basal ganglia (BG) and thalamus. In addition, it also segments the brainstem, the cavum septum pellucidum, the cerebellar vermis, as well as the third and fourth ventricles (A. Uus et al., 2023). We focused our analysis on eight anatomical structures aggregated across hemispheres: eCSF, cGM, WM, LV, cerebellum, BG, thalamus and brainstem, discarding the small cavum septum pellucidum, cerebellar vermis as well as third and fourth ventricles.

### 2.3 Standardized visual quality assessment

We then performed a visual quality assessment following the protocol introduced in Sanchez et al. (2024) and available on Zenodo for reproducibility Bach Cuadra et al. (2025). The visual rating was based on an interface using HTML reports that displayed the data in a standardized layout as illustrated in Figure 1A. The code to generate these reports is publicly available on GitHub. Raters were asked to conduct the quality assessment as described in the protocol and to score the following criteria, as illustrated in Figure 2:

- Is the brain fully reconstructed? [Binary] – Are there holes, or large parts of the brain missing?
- Geometrical artifacts. [Continuous] Did the SRR introduce any non-biological, textured artifact, such as stripes?
- Topological artifacts. [Continuous] Did the SRR introduce any discontinuity in the cortical gray matter (cGM), or are some parts of the white matter (WM) directly touching the cortical cerebrospinal fluid (CSF)?
- Noise. [Continuous] Is there a high level of noise in the image, potential preventing us from seeing the deep gray matter (dGM) clearly?
- Tissue intensity contrast – WM/cGM/cerebrospinal fluid. [Continuous] Is the contrast sufficient? Do we see well the cGM?
- Tissue intensity contrast – WM/dGM. [Continuous] Is the contrast sufficient? Do we see well the sub-regions in the dGM?

**Figure 1:**
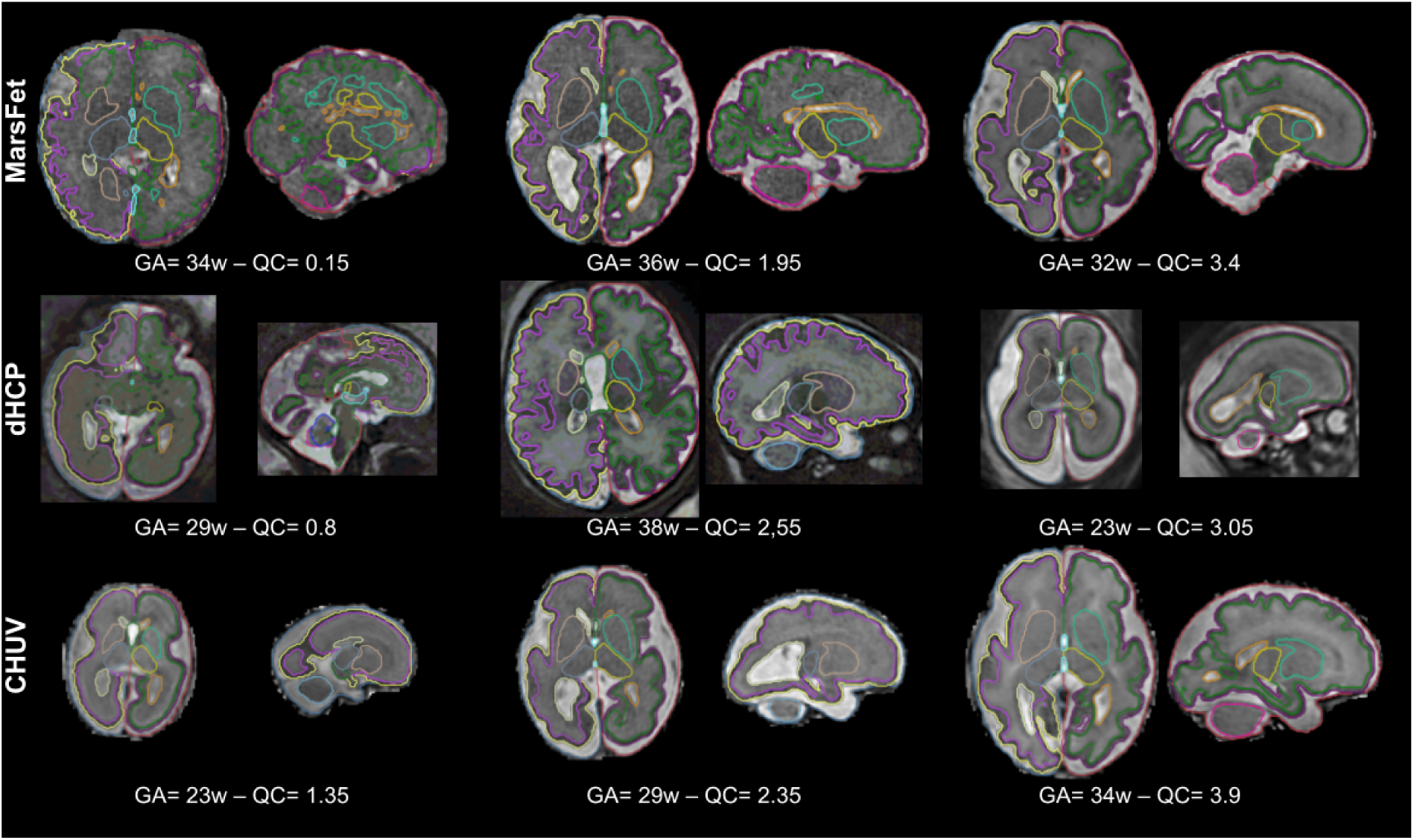
Examples of data from the different cohorts, with different gestational ages and different global QC ratings. Overlaid on each example are segmentations from BOUNTI, before aggregation into the eight categories of interest. Note that two slices are not sufficient to extensively reflect the quality of a given subject, as the rating is based on inspection of *all* slices, and artifacts often lie at the extremes. We refer the reader to Bach Cuadra et al. (2025) for additional examples.

**Figure 2:**
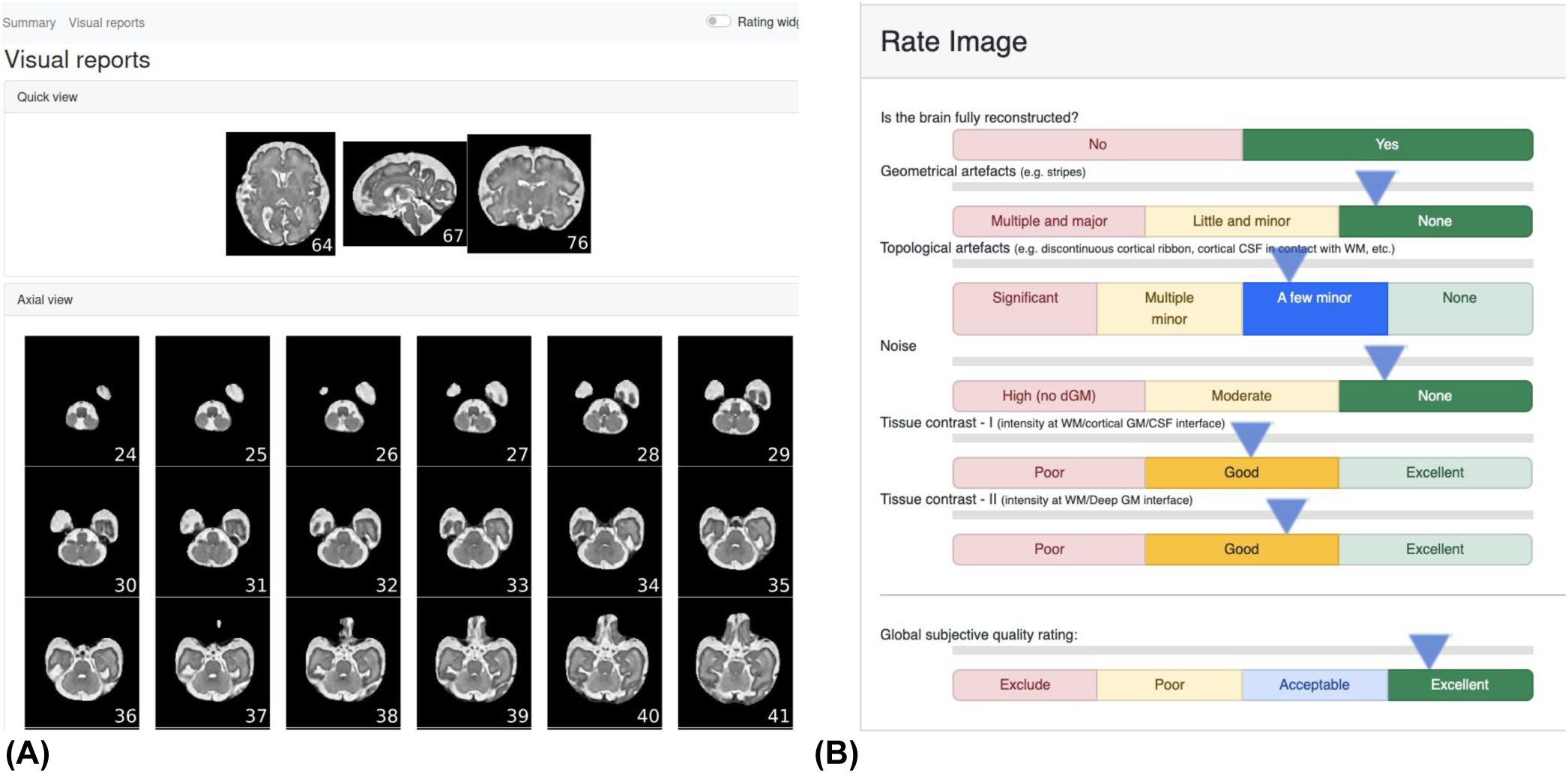
The QC interface used throughout the study. (A) The visualization interface first shows a quick view of each orientation in a given volume, before showing all slices in each orientation. (B) The rating interface, embedded in the HTML report, asks the rater to go through a guided assessment of the quality of the image, by rating 6 different criteria before giving a global subjective quality rating.

All continuous scores range from 0 (very poor) to 4 (excellent), following the scales shown in Figure 2. After scoring the artifacts, the raters were asked to provide a final global subjective quality rating score for the image (on a continuous scale between 0 and 4), based on the previously assessed criteria. This global rating served as our quality reference throughout the study.

#### Rater training

Three raters (AM, GMJ, TS) underwent a training session, where they were asked to rate 25 images from MarsFet. Based on their rating, a discussion session was organized to review cases with strong disagreement. During the discussion, raters discussed extensively the cases they considered as ambiguous to understand the rationale for each rater’s choice and reach a consensus.

After this discussion, each rater evaluated a part of the dataset (AM rated MarsFet + dHCP, GMJ rated BCNatal and TS rated CHUV), resulting in a detailed quality assessment for each of the 635 fetal MRI scans.

### 2.4 Analysis of the impact of image quality on normative models

#### QC Subgroups definition

The distribution of the image quality ratings across the four datasets is shown in Figure 3. Based on this distribution, we defined four quality groups, representing data with increasingly stringent QC criteria:

- All: quality scores ≥ 0 – n=635
- Poor+: quality scores ≥ 1 – n=598
- Accept+: quality scores ≥ 2 – n=485
- GroundTruth: quality scores ≥ 2.7 – n=314

**Figure 3:**
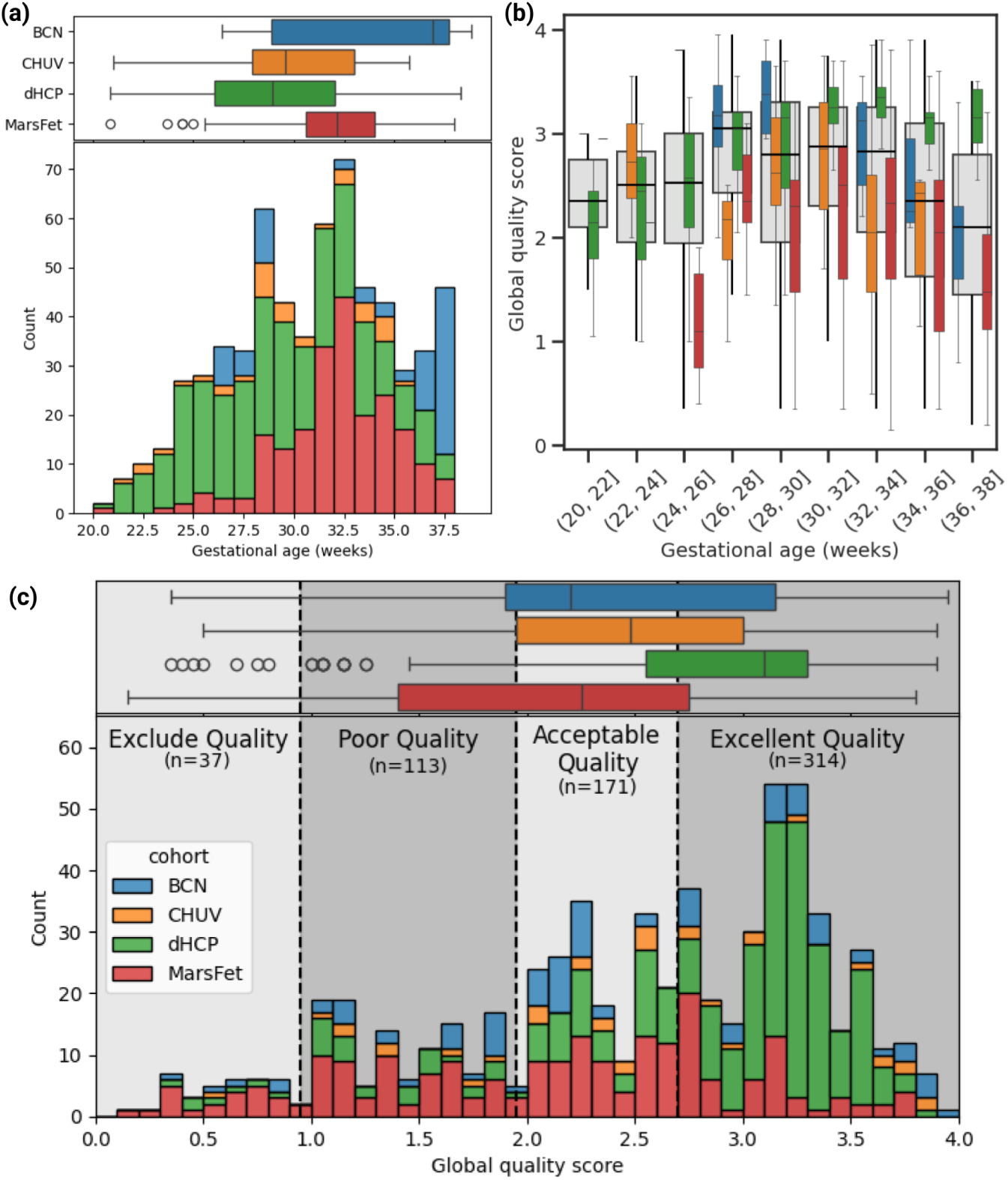
Quality and gestational age (GA) distribution of the data across the different cohorts. (a) Distribution of GA across datasets. (b) Distribution of global quality as a function of GA. (c) Distribution of the global quality score across the different datasets, with the excellent quality defined to include subjects with a score≥ 2.7.

The threshold value of 2.7 for the GroundTruth subgroup was determined based on an experiment described in Supplementary Section S2 and Figure S1. We started from the full cohort of 635 subjects, which was ranked according to QC score. We then iteratively removed the lowest-quality data in batches of ten, generating a series of subgroups with progressively higher data quality and decreasing sample size. For each subgroup, we assessed the stability of the fitted normative models using bootstrapping. As expected, model variance increased markedly when the sample size dropped below approximately 250 subjects (Bozek et al., 2023). A QC threshold of 2.7 was therefore selected, as it ensured stable normative model estimates while retaining a sufficient sample size (n = 314). Visual inspection further confirmed the high quality of all data with QC scores above this threshold.

#### Harmonization on volumetric measurements

After the categories were defined, data from each category were harmonized using the ComBat-GAM method (Pomponio et al., 2020), which extends the original ComBat algorithm (Johnson et al., 2007) by incorporating generalized additive models to flexibly adjust the nonlinear effect of age while removing site-related variance. This model was run by using age as a smoothing term and covarying for total intracranial volume. Harmonization was performed separately for the four nested quality datasets: All, Poor+, Accept+, GroundTruth.

#### Normative models specification and centile trajectories

As outlined by Marquand et al. (2016), the objective of normative modeling extends beyond simply estimating a regression curve that captures the population average. It also involves determining the centiles of the distribution for each brain phenotype, allowing quantification of individual deviations from the reference model. In this study, we employed the generalized additive model for location, scale, and shape (GAMLSS) framework to analyze and characterize how each brain feature, within each QC subgroup, evolves with age as participants progress through gestation. This approach, originally introduced by Stasinopoulos and Rigby (2008), provides a robust and flexible framework for normative modeling. We adopted the implementation of GAMLSS developed by Dinga et al. (2021), which builds upon the R package described in Stasinopoulos and Rigby (2008). This implementation includes automated parameter tuning to optimize model fitting. In our analysis, GAMLSS models were fit using the Sinh–Arcsinh (SHASH) distribution with cubic spline smoothing, and where two parameters out of four were modeled as a function of age, namely location *µ* scale *σ*, Skewness and kurtosis were estimated from the data, but were made independent from gestational age, following Bozek et al. (2023) and Dinga et al. (2021).

Within each QC subgroup, we fit a GAMLSS model and obtained centile trajectories at the 5th, 50th, and 95th percentiles (as well as 1st, 10th, 90th and 99th in some cases). Gestational age was rounded to two decimal places to stabilize fits and align ages across prediction curves. We assessed the stability of the GAMLSS fitting across different population samples via bootstrapping: for each QC subgroup, we fit 50 GAMLSS models from different bootstrap-resampled populations from the previously harmonized data, and reported the average and standard deviation across the centiles predicted from these models at each gestational age.

#### Metrics

To quantify how much a subgroup’s predicted curve at a given centile deviates from the GroundTruth curve at the same centile for a given cortical feature, we computed the mean *E*_1_ error as proposed in (Bozek et al., 2023). The error *E*_1,*p*_(*a*) is defined as the distance between the curves of two GAMLSS models at centile p and given gestational age *a*:

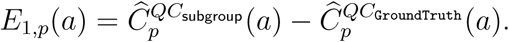

*Ĉp* is the predicted centile for a model, and *QC*_subgroup_ is the given subgroup compared against the GroundTruth subgroup. For each feature×subgroup×centile combination, we summarized a given centile’s deviation (or bias), by computing the E_1_ mean error averaged across gestational ages, namely

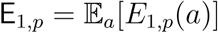

where *a* is sampled on a uniform grid of gestational ages between 20 and 39 weeks.

E_1,*p*_ > 0 indicates that a subgroup’s curve lies above the ground truth’s curve at the same centile (higher predicted values), while an E_1,*p*_ < 0 indicates that a subgroup’s curve lies below that of the ground truth at the same corresponding centile (indicating lower predicted values). Values of E_1,_*_p_* ≈ 0 indicate no systematic bias. As the scale of E_1_ varies with the size of the structure being investigated, we also reported a normalized E**_1_** error:

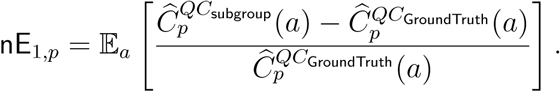

#### Statistical comparisons

Statistical analysis was run on the E_1_ error distribution estimated via the 50 bootstrapped GAMLSS models. For each brain feature, centile and QC subgroups, we tested two hypotheses. First, to assess the presence of bias in a given subgroup, we used a Wilcoxon rank-sum test (non-parametric and non-paired test). Namely, we tested whether the E_1_ error between a given subgroup and the GroundTruth subgroup significantly differed from zero. Secondly, we tested whether increasing the QC stringency (e.g. from All to Accept+) would lead to statistically significant differences in E_1_ errors between the subgroups. To do so, we assessed the difference across QC subgroups using a Wilcoxon signed-rank test.

### 2.5 Disentangling the impact of sample size and image quality

Up to this point, the proposed design did not allow us to separate the impact of data quality from that of sample size on a GAMLSS model, because QC involved removing data points below a given threshold, leading to nested samples of decreasing size as quality thresholding became more stringent (e.g. *n* = 314 for GroundTruth vs *n* = 635 for All).

To disentangle these effects, we needed to control for both the number of data points included in a given sample (n_samp_) and for the data quality of the sample. To achieve this, we implemented a weighted bootstrapping strategy. For a data point *i* with quality score *q_i_*, the resampling probability was defined as

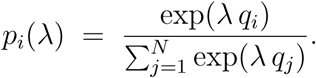

Here, *λ* is a tuning parameter. The expected mean quality of the bootstrap sample was then

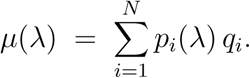

By solving the root equation *µ*(*λ*) = *q*^∗^ for *λ*, where *q*^∗^ is the desired target quality of the sample, the bootstrap resampling scheme yielded (in expectation) samples whose average quality matched the target.

Reweighted bootstrapping allowed us to obtain bootstrap samples of size n_samp_ and average quality *q*. We then defined an equally spaced grid of sample sizes n_samp_ ∈ {200, 350, 500, 650} and quality levels *q* ∈ {1.5, 2.0, 2.5, 3.0}. For each of the sixteen combinations, we bootstrapped 10 samples, each of which was used to fit a GAMLSS model. This procedure was repeated for each brain feature and yielded an array of models fit on data with varying sample sizes and average quality.

We then quantified the impact of n_samp_ and quality on the E_1_ error distributions at the 5th, 50th, and 95th centiles across brain structures. To this end, we fit a linear mixed-effects (LME) model with the following structure:

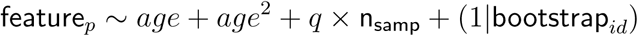

where residuals were assumed to be Gaussian and random intercepts were fit for each bootstrap repetition (bootstrap*_id_*). We also compared this model with an alternative specification in which the variance of the residuals was weighted according to n_samp_, following the hypothesis that estimation variance decreases with increasing sample size. Specifically, we assumed

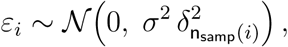

where 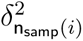 denotes the variance multiplier associated with each n_samp_ level. The significance of the estimated coefficients was assessed using Wald *t*-tests. We report effect sizes (*β* coefficients), *p*-values, and normalized effect sizes. Because the fit coefficients *β* are tied to the scale of the variables they model—expressing average change per unit change in the predictor—and because brain features have markedly different volumes, normalized effect sizes were computed by dividing each coefficient by the average feature volume. Results were corrected for multiple comparisons using Bonferroni correction across centiles.

#### Code

All analyses were implemented in R using gamlss, mgcv, MASS, dplyr, tidyr, nlme, rstatix, and ggplot2; and in Python using pandas, numpy, neuroHarmonize, and matplotlib. Data tables (CSV files) and the code used to analyze and visualize the data will be released upon acceptance of the paper.

## 3 Results

For the sake of space and clarity, we report the main results using three structures of interest that are representative of the results obtained on the other structures: cerebral white matter, lateral venticles and cerebellum. We report the results for the other structures as Supplementary Material.

### 3.1 Harmonization can obfuscate QC-related effects

We first examine the interaction between QC and harmonization in the context of normative modeling. Figure 4 presents scatter plots of raw volume data (without harmonization) as a function of gestational age for three representative structures, with QC scores indicated by color. The right column shows detrended plots (subtracting the average trajectory) alongside the average quality marginalized across detrended volumes.

**Figure 4:**
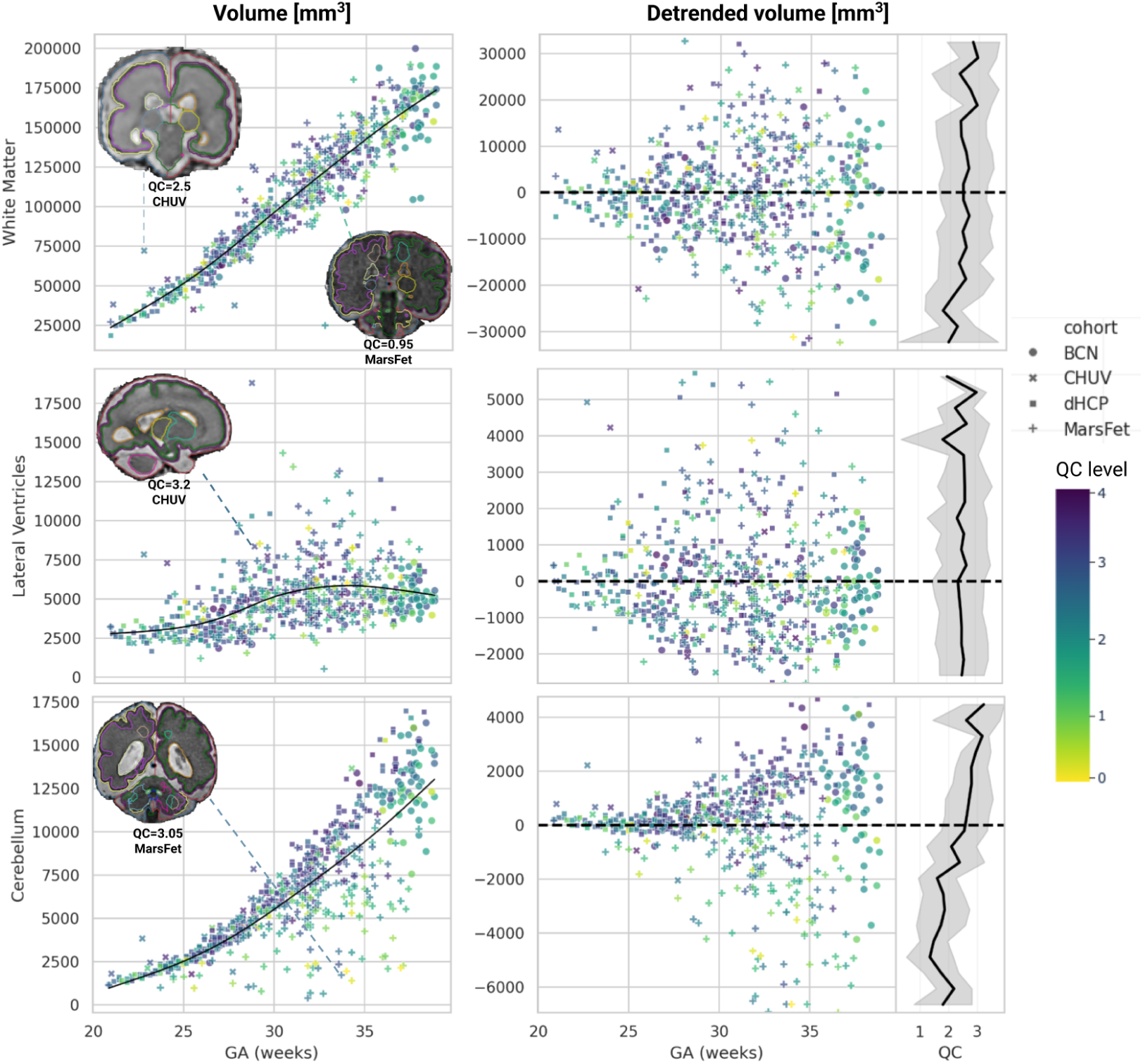
Influence of QC on raw data. *(Left)* Volume as a function of gestational age for raw volumes (not harmonized), for the three structures of interest. The color of the points encodes the QC level. The black line is a spline fit on the data. *(Middle)* Detrended volume, with the spline average trend subtracted from the original volume data. *(Right)* Average data quality as marginalized across the y-axis of the detrended volume plot.

This visualization clarifies that there is no simple relationship between QC scores and distance to the mean trajectory (residuals from regression). Poor quality images do not systematically produce outlying data, and conversely, high quality images can correspond to data samples located relatively far from the mean. Moreover, we observe marked variation across structures in how QC impacts the data distribution relative to gestational age. While lower QC values visibly associate with reduced cerebellar volumes, this pattern does not hold for lateral ventricles or white matter—particularly evident in the detrended plots (right column). To confirm this absence of a straightforward relationship, we computed pairwise correlations between detrended volume and QC, obtaining consistently low explained variance (*R*^2^ < 0.05) across all structures. Supplementary Figure S4 confirms these observations for the remaining structures.

We then examine the critical interaction between QC and harmonization. Figure 5 shows normative curves fit on all data (top row) versus the GroundTruth subgroup (bottom row), without (left column) and with (right column) harmonization, for the cerebellum. As expected, harmonization clearly reduces data spread around the median trajectory and yields centile curves closer to the median. This effect is more pronounced for all data (top row) than for the GroundTruth subgroup (bottom row), since the former includes poor quality measurements located far from the median trajectory (top left plot). Critically, when harmonization is applied in the presence of poor quality measures (top row), the QC-related spread is compensated such that many poor measurements shift close to the median trajectory (top right), specifically within the 5th–95th centile curves typically used to identify abnormal data in normative models. Similar patterns emerge across other structures (Supplementary Figures S2 and S3).

**Figure 5:**
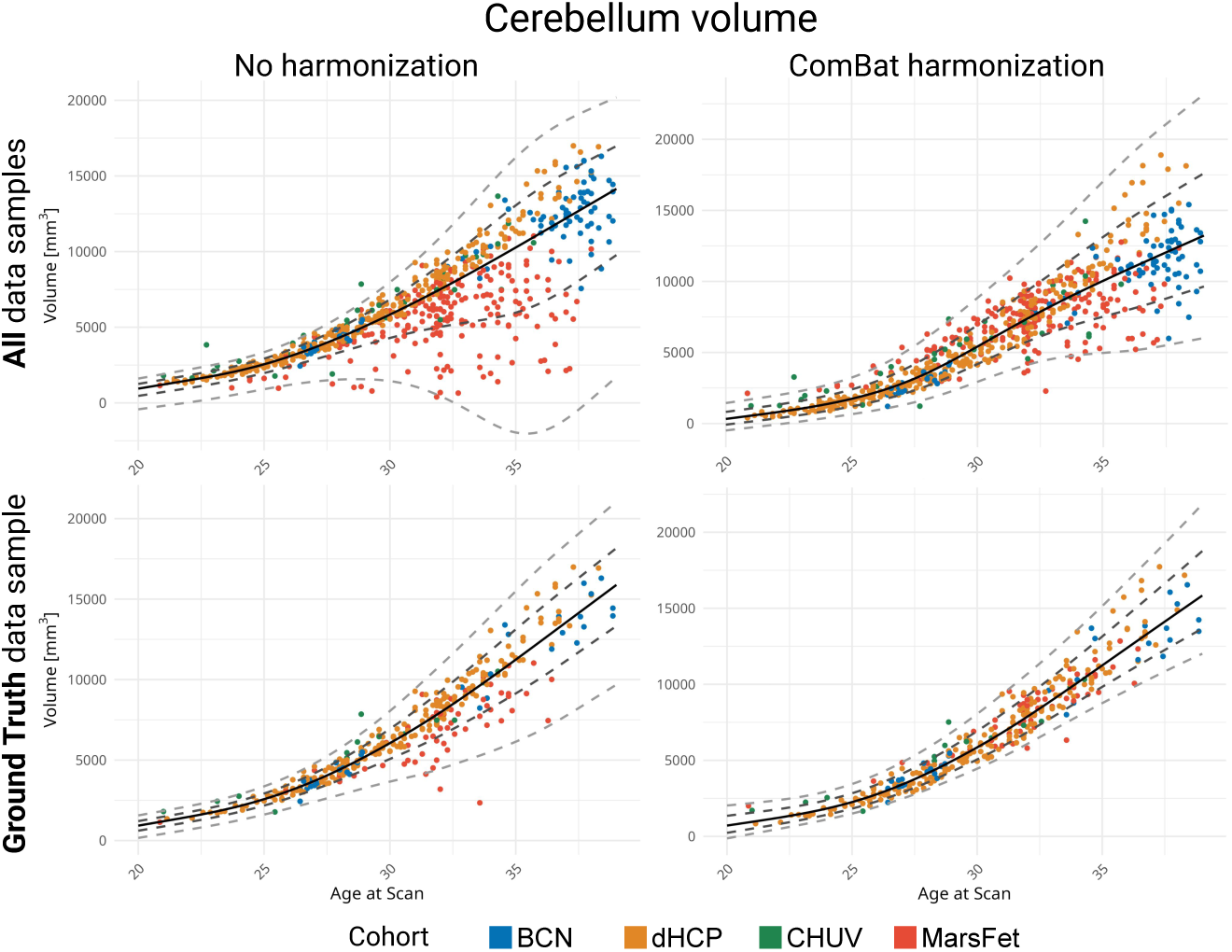
Interaction between QC and harmonization. Normative curves at centiles 1st, 5th, 50th, 95th and 99th, for the cerebellum, fit on all data (top row), or on the GroundTruthsubgroup (bottom row), and featuring raw (left) or harmonized (right) data.

These results illustrate the substantial impact of QC on normative curves and its non-trivial interaction with harmonization.

### 3.2 Poor quality images can bias normative trajectories

Figure 6 illustrates the impact of image quality on the normative curves for white matter, lateral ventricles and cerebellum volume. The results for the remaining structures are shown in Supplementary Figures S6 and S5. The first observation from Figure 6 is the progressive impact of image quality on centile curves: deviations from the GroundTruth subgroup increase as lower-quality data are included (Accept+< Poor+< All). The second striking observation is that the impact of image quality on the normative trajectories varies across structures.

**Figure 6:**
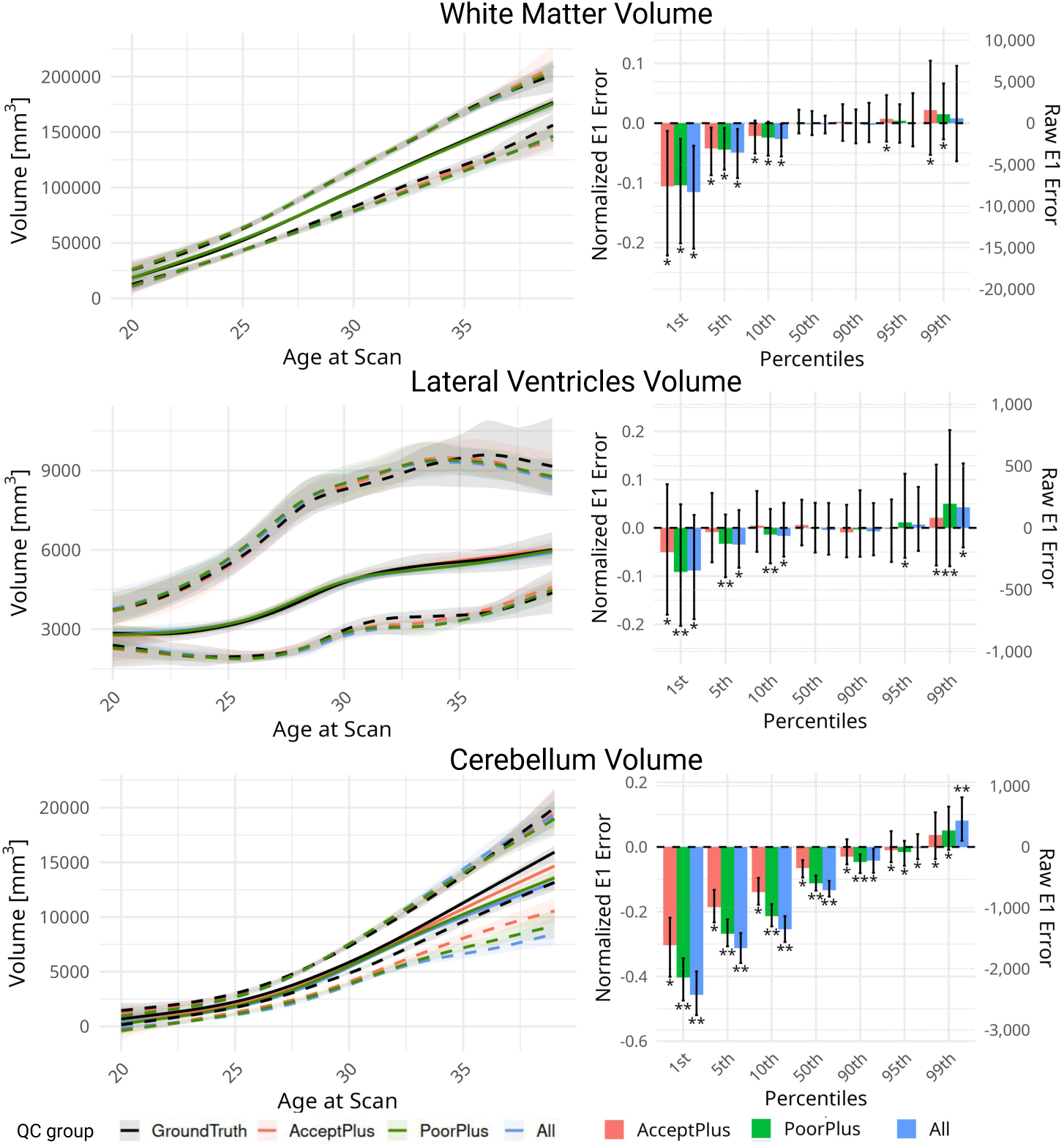
Impact of data quality on normative curves. Left column: Bootstrapped curves with 95% confidence intervals for the 5th, 50th and 95th centiles. Right column: E_1_ error distribution with 95% confidence intervals computed for 1st, 5th, 10th, 50th, 90th, 95th and 99th centiles. A single ⭐ indicates that the mean is significantly different from 0, whereas ⭐⭐ indicates a significant difference from 0 and from the other quality subgroups.

For the white matter volume, the impact of image quality on the median trajectory (50th centile) is limited, and QC biases more strongly the lower centiles (1st, 5th, and 10th) than the higher centiles (90th, 95th, and 99th). For the lateral ventricles volume, the confidence intervals around the centile curves are substantially wider, indicating greater variability across bootstrap samples compared with white matter. As for white matter, the effect on the median trajectory is small, but increases for more extreme centiles. In contrast, cerebellar volume shows an asymmetric effect of image quality relative to the median trajectory. Lower centiles exhibit a strong negative bias, whereas higher centiles are positively affected. Notably, the effect remains negative and pronounced for the median trajectory. Overall, these results indicate that the impact of image quality is not uniformly distributed around the median: the inclusion of lower-quality data introduces a systematic bias, manifesting as a downward shift affecting both lower centiles and the median trajectory.

Observations for the remaining structures (Figures S6 and S5) are consistent with these patterns. The effect of QC resembles that observed for white matter in cortical gray matter, basal ganglia, and thalamic volumes; it is similar to that of lateral ventricles for eCSF, and comparable to cerebellar volume for the brainstem.

### 3.3 The observed bias is not due to variations in sample size

To confirm that data quality is the main factor inducing the observed bias, and confirm that it is not a consequence of an interaction with the number of samples used to fit the normative model, we carried out an experiment using weighted bootstrapping to build populations of size n_samp_ ∈ {200, 350, 500, 65} and quality *q* ∈ {1.5, 2.0, 2.5, 3.0}. For each brain structure, and (n_samp_, *q*) combination, we generated 10 bootstrap samples used to fit different normative models. Using a LME model, we were then able to disentangle the effect of n_samp_ and *q*.

Results on Figure 7 show clearly that the bias affecting the centile curves is consistent across n_samp_. This is confirmed in the right-most column of the figure, where we superimpose the curves obtained with different n_samp_ for a given QC subgroup *q*. The observations for the other brain structures(Fig. S7) are consistent.

**Figure 7:**
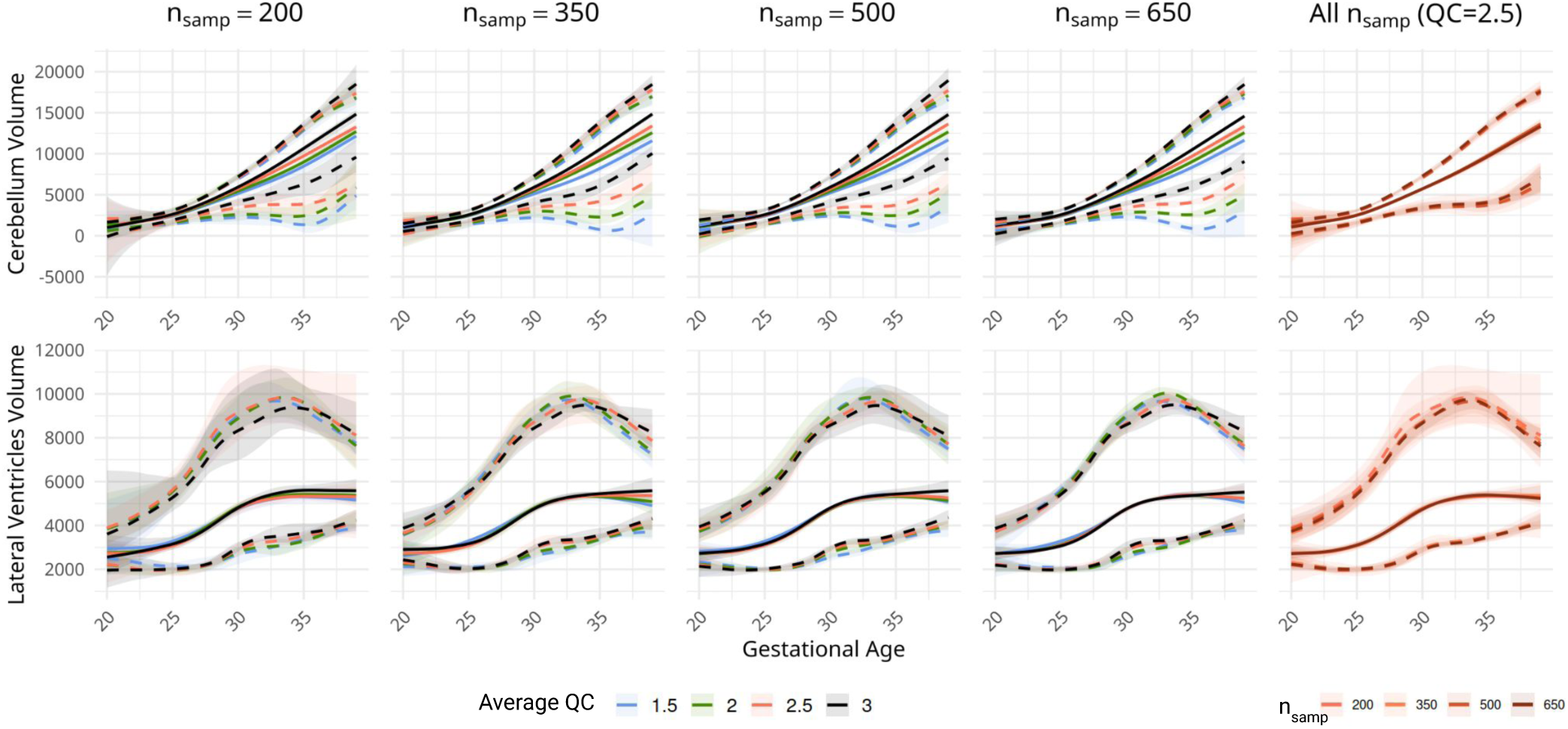
Bootstrapping samples confirm a consistent bias induced by QC. Weighted bootstrapping allows obtaining samples with a controlled n_samp_ as well as *q*. The first four columns show normative centile curves at various n_samp_ (200, 350, 500, 650) for different *q* (1.5, 2.0, 2.5, 3.0), and the last column shows all n_samp_ for a fixed quality (2.5).

Quantitative results from the LME analysis are presented in Table 2 (with its extended version shown in Table S2). The results indicate that, except for the lateral ventricles, there is a consistently significant effect of QC on the estimated centiles. The effect size is orders of magnitude larger than the effect size of n_samp_. Note that this difference is (partially) offset by the difference in scale between the variables.

**Table 2:** Effect size from a LME model shows that the impact of QC on estimated centiles is consistent, large, and asymmetrical. Grayed-out cells represent non statistically significant results, raw effect sizes are the slopes (*β*) of the fit LME model.

| Region | Centile | QC effect size |  | n<sub>samp</sub> effect size |  | Interaction<br>Eff. size |
| --- | --- | --- | --- | --- | --- | --- |
|  |  | Raw | Relative | Raw | Relative |  |
| eCSF | 5th | 5574.99 | 0.11 |  |  | -1.18 |
| | 50th | 4807.19 | 0.068 | 1.85 | $2.63 \times 10^{-5}$ | -0.82 |
|  | 95th | 4475.48 | 0.048 |  |  |  |
| Cortical Gray Matter | 5th | 2140.03 | 0.058 | 1.79 | $4.88 \times 10^{-5}$ | -0.85 |
| | 50th | 688.15 | 0.015 | -1.12 | $-2.42 \times 10^{-5}$ | |
| | 95th | 858.24 | 0.015 | -1.88 | $-3.26 \times 10^{-5}$ | 1.24 |
| White Matter | 5th | 4629.56 | 0.062 | 5.09 | $6.77 \times 10^{-5}$ | -3.24 |
| | 50th | 414.67 | 0.004 | -1.26 | $-1.35 \times 10^{-5}$ | 0.48 |
|  | 95th | 1529.11 | 0.014 |  |  |  |
| Lateral Ventricles | 5th | 156.70 | 0.057 | 0.20 | $7.19 \times 10^{-5}$ | -0.08 |
|  | 50th |  |  |  |  |  |
| | 95th | | | 0.53 | $7.30 \times 10^{-5}$ | -0.15 |
| Cerebellum | 5th | 1528.81 | 0.57 |  |  |  |
| | 50th | 616.71 | 0.106 | -0.49 | $-8.31 \times 10^{-5}$ | 0.16 |
| | 95th | 402.40 | 0.052 | 0.46 | $5.90 \times 10^{-5}$ | |
| Basal Ganglia | 5th | 148.48 | 0.040 |  |  |  |
| | 50th | -51.95 | -0.011 | -0.13 | $-2.84 \times 10^{-5}$ | 0.06 |
| | 95th | -460.19 | -0.073 | 0.38 | $5.99 \times 10^{-5}$ | -0.12 |
| Brainstem | 5th | 525.17 | 0.208 |  |  |  |
| | 50th | 169.06 | 0.047 | -0.17 | $-4.70 \times 10^{-5}$ | 0.05 |
| | 95th | 66.67 | 0.015 | 0.13 | $2.97 \times 10^{-5}$ | |
| Thalamus | 5th | 105.13 | 0.035 | -0.07 | $-2.38 \times 10^{-5}$ | 0.03 |
|  | 50th | 15.92 | 0.004 |  |  |  |
|  | 95th | -121.26 | -0.026 |  |  |  |

To contextualize these effect sizes, consider that in our experimental design, quality levels differ by 0.5 units (e.g., from 2.0 to 2.5) while sample sizes differ by 150 subjects (e.g., from 200 to 350). For eCSF’s 50th centile, a 0.5-unit increase in average QC corresponds to a 3.4% volume increase (0.068 × 0.5), whereas adding 150 subjects yields only a 0.4% increase (2.63 × 10^−5^ × 150). This illustrates how QC has a substantially larger impact than sample size on the observed biases.

These effects are not mitigated with the interaction between the variables, that is generally much smaller compared to the effect of QC (rightmost column). This further confirms that data quality biases normative centile curves: irrespective of the amount of data used to fit a model, the average data quality has a systematic and consistent biasing effect. Furthermore, except for the basal ganglia and thalamus, this effect shows an asymmetric and non-uniform downward bias induced by lower quality data: including lower quality data generally draws the estimated volumes down, and has a stronger effect on inferior centiles compared to upper ones.

## 4 Discussion

### Quality-induced bias in the context of early brain development

Previous studies focusing on QC have demonstrated that poor data quality (often driven by motion) biases image-derived biomarkers (Backhausen et al., 2016; Gilmore et al., 2021; Reuter et al., 2015). However, these analyses are typically static, comparing adult groups without explicitly accounting for developmental trajectories. In fetal imaging, such an approach is not relevant, as rapid brain development necessitates explicit modeling of gestational age. By incorporating a temporal dimension through normative modeling, our results extend prior findings to the fetal domain and demonstrate that data quality biases not only individual biomarkers but also the estimated developmental centiles themselves.

### Data quality can bias normative models

Our results demonstrate the critical influence of data quality on normative modeling. Including lower-quality data systematically biases estimated centiles, with the largest effects observed at the distribution extremes, particularly in the lower centiles (1st–10th). Importantly, these effects do not arise from a uniform increase in variance: poor-quality data can bias normative curves asymmetrically across centiles, and may even affect the median trajectory. Failing to exclude poor-quality data prior to fitting normative models therefore leads to systematic, non-trivial bias patterns in the resulting centile curves.

These findings are particularly important from a clinical perspective, as centiles are often the most actionable outputs of normative models, informing the detection of atypical developmental trajectories and clinical risk stratification (Marquand et al., 2016). This susceptibility raises concerns regarding the clinical reliability of normative models derived from datasets with insufficiently well-defined QC procedures.

Although prior fetal normative studies generally report having performed QC, the absence of clearly defined and reproducible QC protocols makes it difficult to evaluate the extent to which their reported centiles may be biased. Importantly, by leveraging a continuous QC scale, we show that bias increases progressively as QC criteria are relaxed, rather than emerging only from severely degraded or obviously corrupted scans. This continuum challenges binary inclusion–exclusion approaches to QC and highlights the need for more nuanced strategies, as even moderate reductions in QC stringency can lead to systematic shifts in centile estimates.

Taken together, these results underscore the necessity of adopting standardized and transparent QC protocols at the community level. Such efforts will be essential not only to improve reproducibility and consistency across normative modeling studies, but also to support the safe translation of these models into clinical practice.

### Bias–variance tradeoff and implications for fetal normative modeling

Our results highlight a fundamental bias–variance tradeoff in normative modeling. Recent work on normative modeling has emphasized the importance of large sample sizes for estimating reliable developmental trajectories (Bozek et al., 2023). Our results add an important nuance to this message: while sample size matters, data quality is a critical determinant of bias in normative centiles, that cannot be compensated by increasing the sample size. Using a larger number of lower-quality scans leads to centile estimates with lower variance but increased bias, whereas restricting analyses to higher-quality data reduces bias at the cost of increased variance due to smaller effective sample sizes. This tradeoff is particularly important in fetal neuroimaging, where sample sizes are typically small (n=127 in (Kyriakopoulou et al., 2017); n=122 in (Machado-Rivas et al., 2022); n=340 in (X. Xu et al., 2024)). In this context, increasing sample size at the expense of data quality could paradoxically degrade the reliability of derived growth trajectories and compromise their clinical utility. Our bootstrapping experiments highlighted a dramatic increase of variance when fewer than n=250 samples were used to fit a normative model.

### Interactions between image quality and segmentation accuracy

An important observation from our analyses is that, although image quality clearly biases normative centiles, disentangling its effects from those related to segmentation accuracy remains challenging. Many scans rated as lower quality do not exhibit overt segmentation failures (Figures 1 and 4), yet their inclusion still leads to systematic shifts in centile estimates. Conversely, some scans with apparently good image quality yield poor segmentations for specific structures, as illustrated by segmentation failures for the cerebellum in a subset of the MarsFet dataset.

While it is common—even in adult neuroimaging studies—to rely exclusively on image-level QC (Backhausen et al., 2016; Esteban et al., 2017; Reuter et al., 2015; Rosen et al., 2018), the most effective QC strategy should both integrate image-based and segmentation-based quality criteria (Klapwijk et al., 2019; Monereo-Sánchez et al., 2021). Implementing such approaches is particularly challenging in the fetal imaging context, however, as there is currently no widely adopted segmentation framework, like FreeSurfer (Fischl, 2012), that could serve as a common reference standard. In the absence of such consensus, we encourage researchers developing new segmentation methods to report failure modes in a systematic and detailed manner. Such transparency would be instrumental in enabling end users to design effective QC protocols, and in accounting for the complex interplay between image and segmentation quality in downstream analyses.

### Limitations and future works

This work currently has some limitations. First, our QC protocol relied exclusively on image-level quality assessment and did not incorporate segmentation-level QC metrics. An important future research direction should be the design and release of a standardized QC protocol to assess the quality of the segmentation, pursuing the efforts initiated by Bach Cuadra et al. (2025).

Second, while we relied on BOUNTI (A. U. Uus et al., 2023) for the segmentation in this work, this algorithm was only designed to handle healthy subjects. There is then a great need to develop tools able to segment both healthy and pathological subjects, as the usefulness of normative models will be in their ability to correctly flag pathological subjects as outliers, which supposes first a correct segmentation of such subjects.

Third, although our study includes the largest multi-centric fetal brain cohort to date (n=635), this sample size remains below the recommendations for normative modeling (Bozek et al., 2023). As such, the precision of the estimation of the outer centiles might still be limited by the sample size.

Fourth, our analyses focused primarily on typically developing fetuses. While this was necessary to establish normative trajectories, the impact of quality-induced bias on pathological cases, where accurate centile placement is most critical, remains to be fully characterized.

Finally, although we relied on state-of-the-art methods for the processing of our data (Sanchez et al., 2025), the generalizability of our findings to imaging protocols beyond fast spin echo and to other segmentation pipelines (Zalevskyi et al., 2024) should be investigated in future studies.

## 5 Conclusion

In conclusion, our results demonstrate that data quality is a major and often underappreciated source of bias in fetal brain normative modeling. While increasing sample size is commonly viewed as the primary avenue to improve the reliability of normative models, our results show that doing so at the expense of data quality can systematically distort centile estimates, particularly at the clinically relevant extremes. Quality-induced bias is subtle, structure-dependent, and progressive, challenging the prevailing view of QC as a simple inclusion–exclusion procedure. Instead, shifting toward a continuous, quantitative characterization of data quality would provide a more nuanced perspective, and enable the discussion of community-accepted levels of QC stringency. Such transparent and standardized QC practices will be essential to ensure the robustness, comparability, and ultimately the clinical utility of fetal brain growth models.

## Author Contributions

*Conceptualization:* AM, TS, GA, MBC. *Methodology:* AM, TS, GA. *Software:* AM, TS, GMJ. *Validation:* AM, TS, GA, MBC. *Formal analysis:* AM, TS. *Investigation:* AM, TS, GMJ. *Data Curation:* NG, MM, EE, VD, MK, LP, JS, AM, TS, GMJ. *Writing - Original Draft:* AM, TS, GA, MBC. *Writing - Review & Editing:* AM, TS, GA, MBC, VD, GP. *Visualization:* AM, TS, GA, GMJ. *Supervision:* GA, MBC, GP. *Project administration:* GA. *Funding acquisition:* GA, MBC, GP, OC, MAGB, EE.

## Data and Code Availability

Tabular data—estimated volumes data and corresponding data quality for each subject—and code will be made publicly available on Zenodo and GitHub upon acceptance of the paper. Sharing of the reconstructed images and segmentations requires data transfer agreements with each of the centers.

## AI use

During the preparation of this work, the authors used ChatGPT and Claude to assist with spell checking, text editing, and language clarity improvements. After using these tools, the authors reviewed and edited the content as needed and take full responsibility for the content of the publication.

## Supplementary Material

### S1 Inclusion and exclusion criteria

To establish a healthy fetal cohort, a multidisciplinary team—including specialists in neuroradiology, obstetrics, and pediatric neurology—developed exclusion criteria to ensure clinical and anatomical normality and standardization across sites. The resulting inclusion and exclusion criteria are detailed in Table S1 below.

**Table S1:**
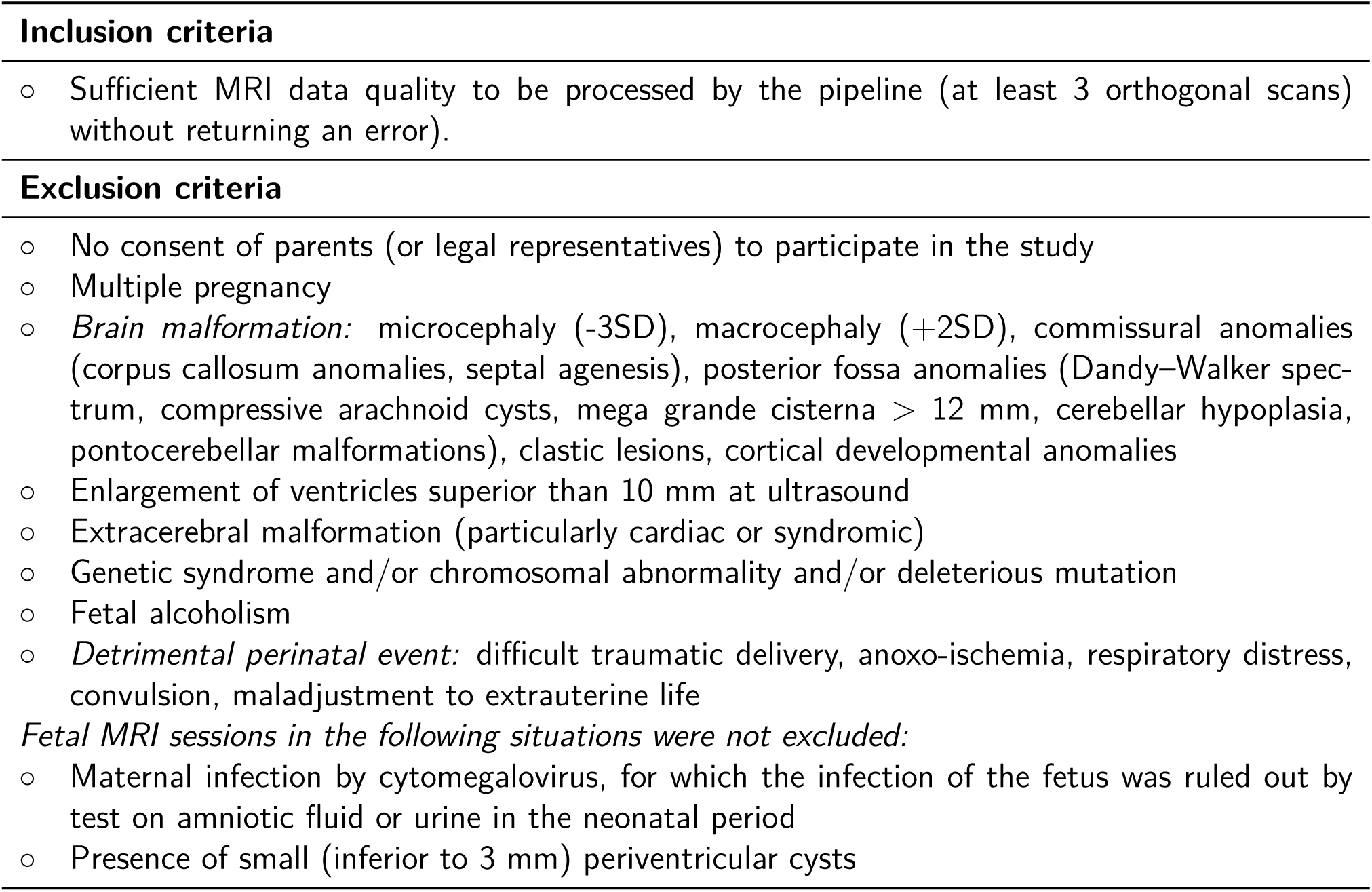
Inclusion and exclusion criteria for the study subjects.

|  |
| --- |
| **Inclusion criteria** |
| ○ Sufficient MRI data quality to be processed by the pipeline (at least 3 orthogonal scans) without returning an error). |
| **Exclusion criteria** |
| ○ No consent of parents (or legal representatives) to participate in the study<br>○ Multiple pregnancy<br>○ *Brain malformation*: microcephaly (-3SD), macrocephaly (+2SD), commissural anomalies (corpus callosum anomalies, septal agenesis), posterior fossa anomalies (Dandy–Walker spectrum, compressive arachnoid cysts, mega grande cisterna > 12 mm, cerebellar hypoplasia, pontocerebellar malformations), clastic lesions, cortical developmental anomalies<br>○ Enlargement of ventricles superior than 10 mm at ultrasound<br>○ Extracerebral malformation (particularly cardiac or syndromic)<br>○ Genetic syndrome and/or chromosomal abnormality and/or deleterious mutation<br>○ Fetal alcoholism<br>○ *Detrimental perinatal event*: difficult traumatic delivery, anoxo-ischemia, respiratory distress, convulsion, maladjustment to extrauterine life<br>*Fetal MRI sessions in the following situations were not excluded:*<br>○ Maternal infection by cytomegalovirus, for which the infection of the fetus was ruled out by test on amniotic fluid or urine in the neonatal period<br>○ Presence of small (inferior to 3 mm) periventricular cysts |

### S2 Iterative subject removal

For this experiment, reported in Figure S1 we started from the entire pool of 635 subjects. We then sorted the data according to QC and iteratively removed the data of lowest quality, 5 by 5 – quality is illustrated on the bottom right of Figure S1. The data were then harmonized on the remaining samples, and then 10 bootstrap samples were used to estimate the variance due to population sampling on the fit GAMLSS models. The normalized E1 error was then computed between the curve obtained at *n*_samp_ and the original curve fit using all subjects.

The results, reported for the 5th and the 50th centile provide a finer version of Figures 6, S6 and S5, showing that a systematic different starts appearing when samples of lower quality are removed. The results also highlight an instability in the model fit when going to *n*_samp_ < 250, with a drastic increase in variance across models. Settling for the ground truth criterion at *QC* = 2.7 allows us to remain far from this high variance zone.

**Figure S1:**
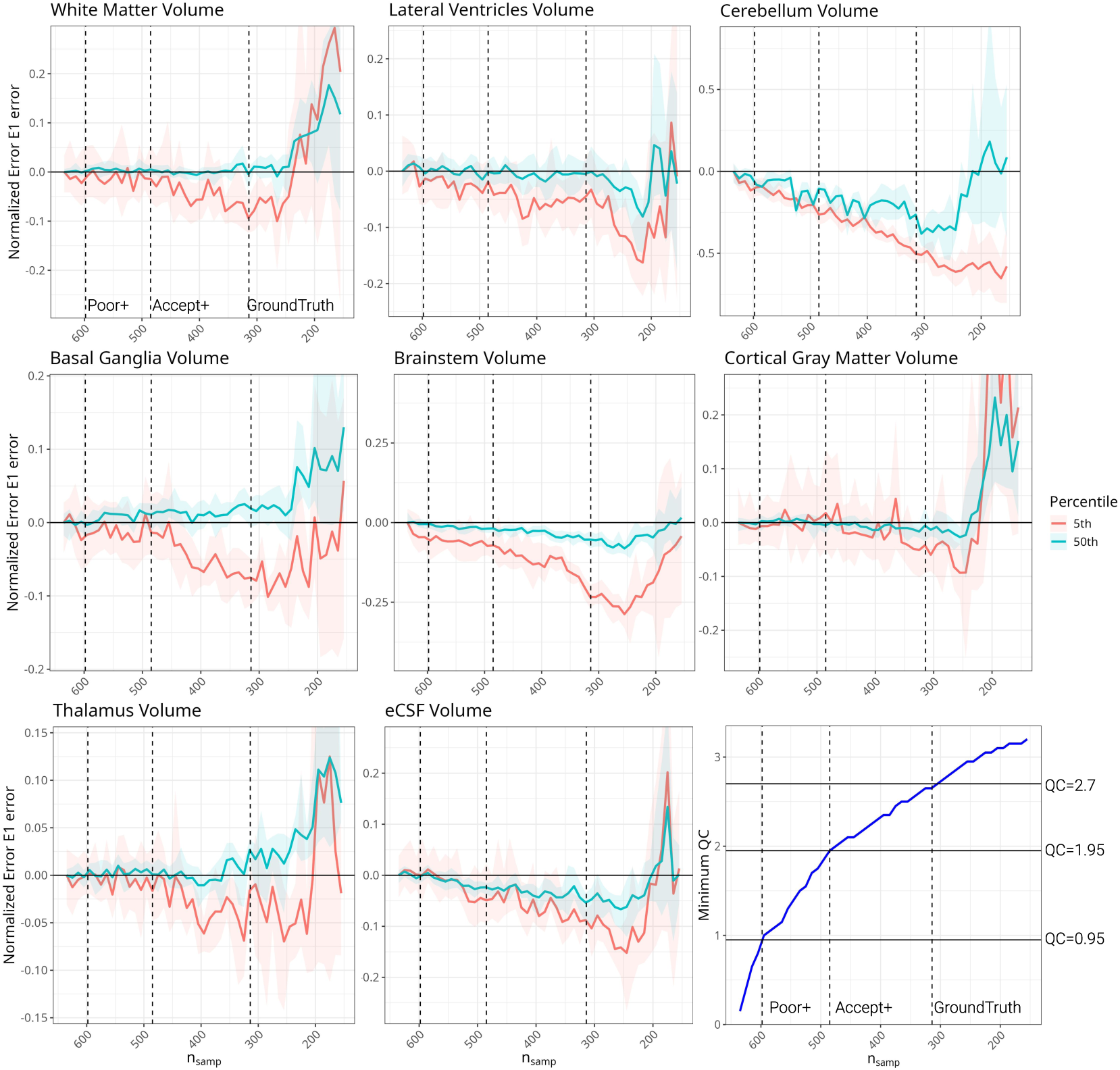
Normalized E1 error against *n*_samp_ < 250 when iteratively removing subjects of low QC values. Thresholds denote the different levels of inclusion criteria discussed in the paper.

### S3 Impact of harmonization on additional structures

Figures S2 and S3 complement the results of Figure 5 and shows how QC and harmonization interact in non-trivial ways. Although the observed effect is generally less dramatic than what was highlighted for the cerebellum, we still observe large differences in predicted centiles, especially for the brainstem, where centiles remain spread independently of harmonization when data of poor QC levels are included in the fit.

**Figure S2:**
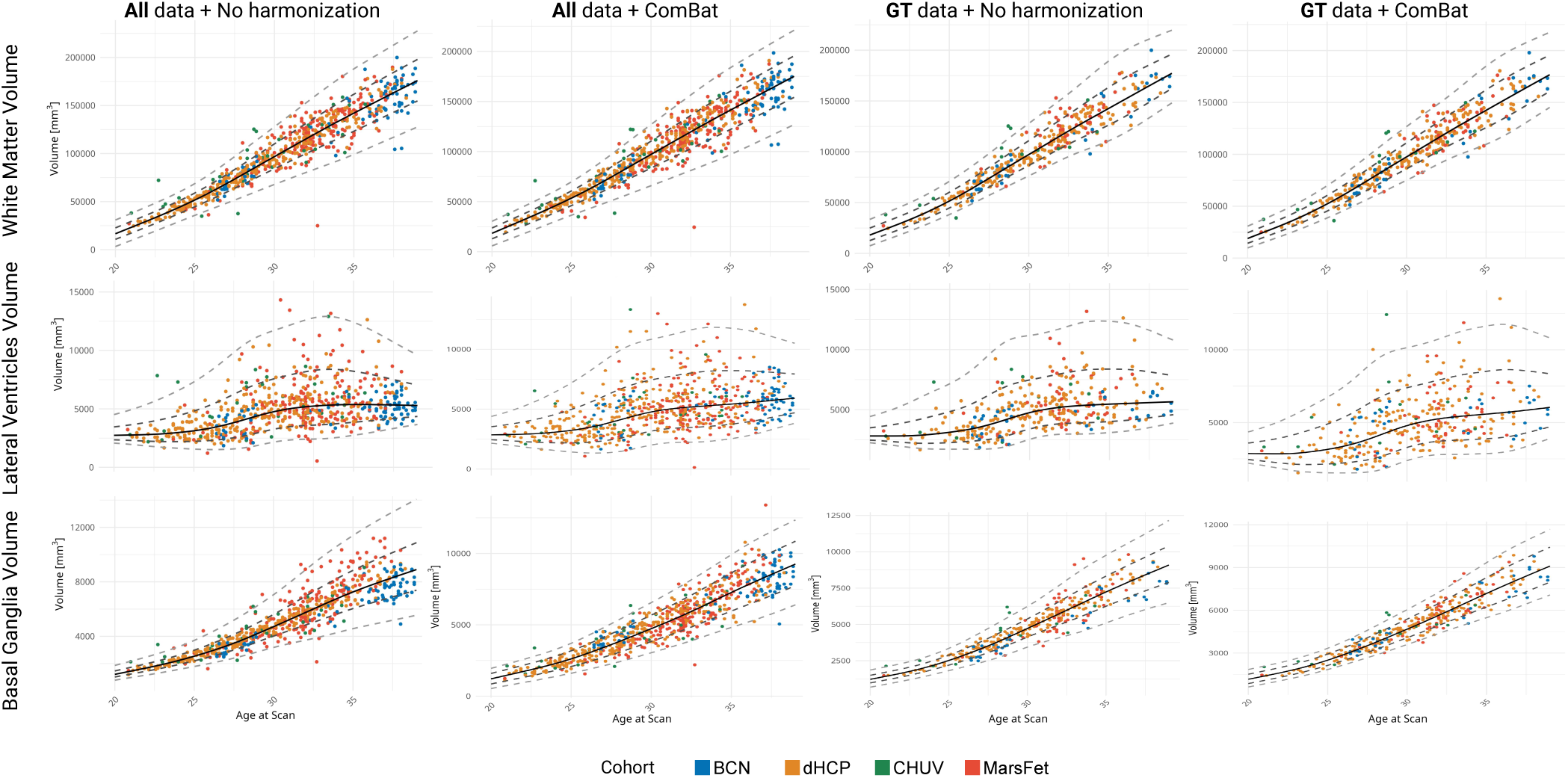
Normative curves at various centiles (1st, 5th, 50th, 95th and 99th) for the white matter (top row), the lateral ventricles (middle) and the basal ganglias (bottom), fit using either all data or QC-ed data (bottom row), and featuring raw or harmonized data.

**Figure S3:**
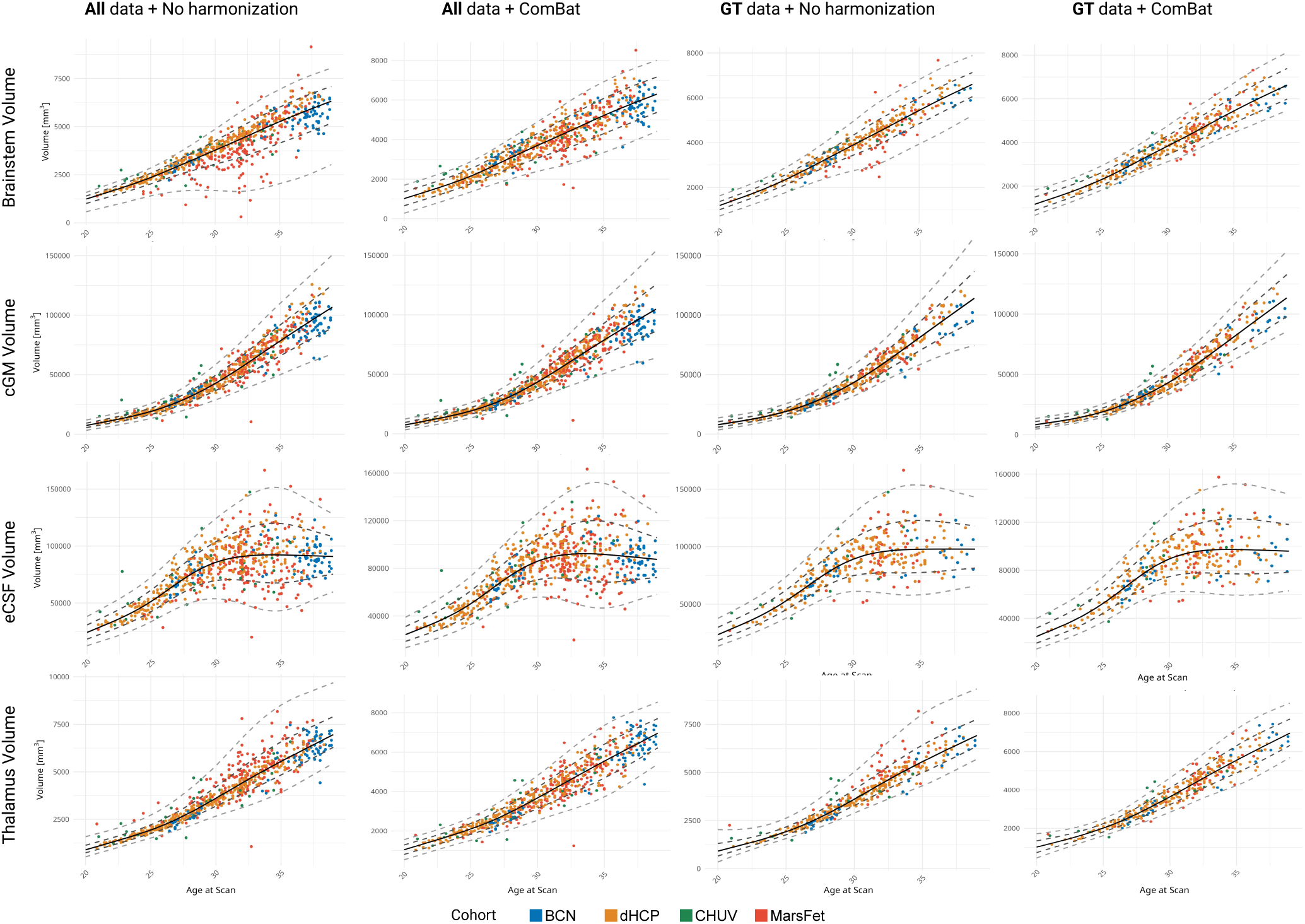
Normative curves at various centiles (1st, 5th, 50th, 95th and 99th) for the brainstem (first row), the cortical gray matter ventricles (second row), the extra-cerebral CSF (third row) and the thalamus (fourth row), fit using either all data or QC-ed data (bottom row), and featuring raw or harmonized data.

### S4 Detrended plots on additional structures

Figure S4 complements Figure 4 and shows globally similar trends. When detrended, there is no obvious connection between outliers and QC.

**Figure S4:**
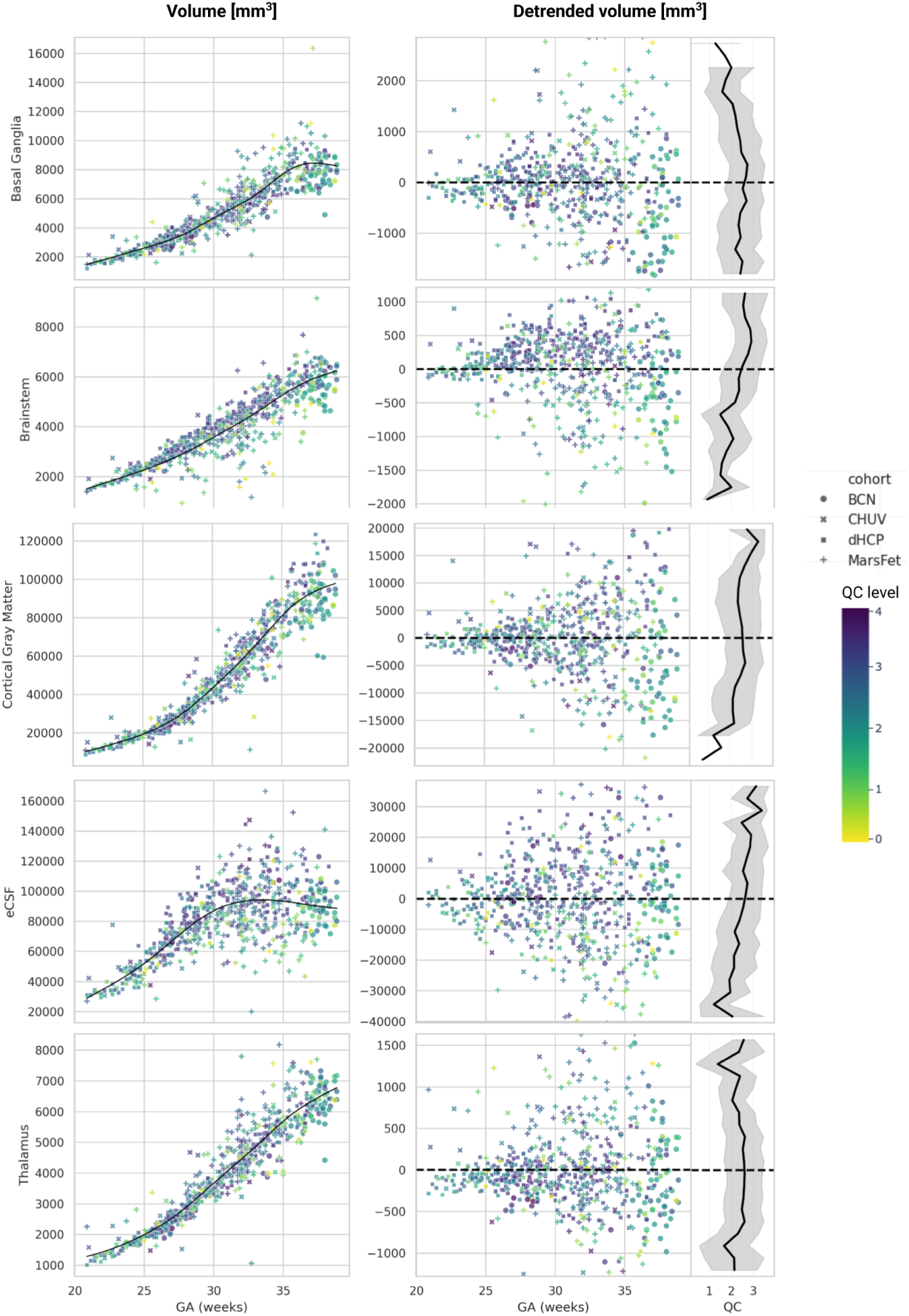
There is no obvious influence of data quality on volumetric estimates. *(Left)* Volume as a function of gestational age for raw volumes (not harmonized). The black line is a spline fit on the data. *(Middle)* Detrended volume, with the spline average trend subtracted from the original volume data. *(Right)* Average data quality as marginalized across the y-axis of the detrended volume plot.

### S5 Normative curves and E1 errors on additional structures

Figures S5 and S6 complement Figure 6 and confirm the strong effect of QC on biasing normative centiles across all brain structures considered.

**Figure S5:**
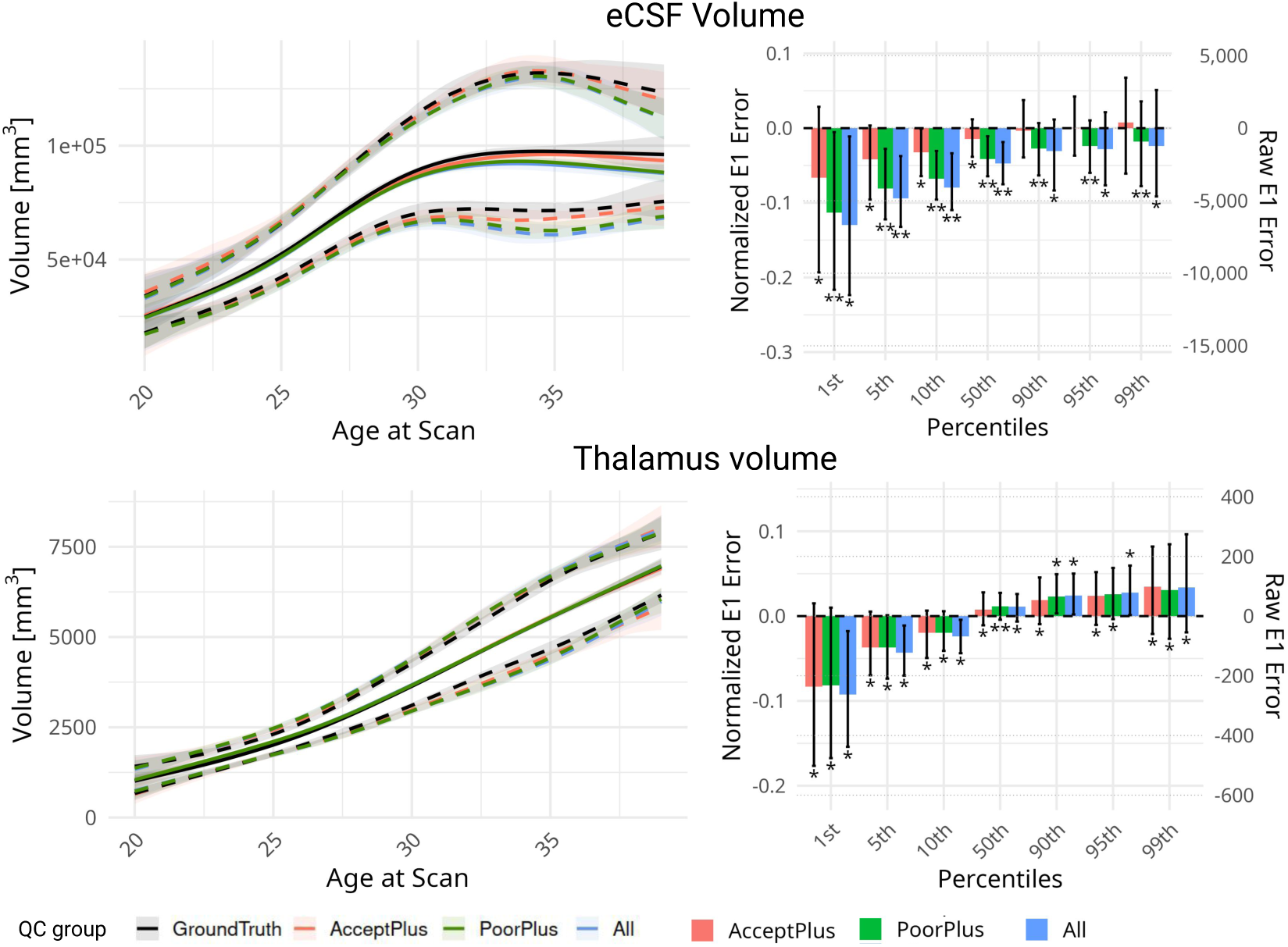
Impact of data quality on normative curves for eCSF, and thalamus (from top to bottom). Left column: Bootstrapped curves with 95% confidence intervals for the 5th, 50th and 95th centiles. Right column: E_1_ error distribution with 95% confidence intervals computed for 1st, 5th, 10th, 50th, 90th, 95th and 99th centiles. A single ⭐ indicates that the mean is significantly different from 0, whereas ⭐⭐ indicates a significant difference from 0 and from the other quality subgroups

**Figure S6:**
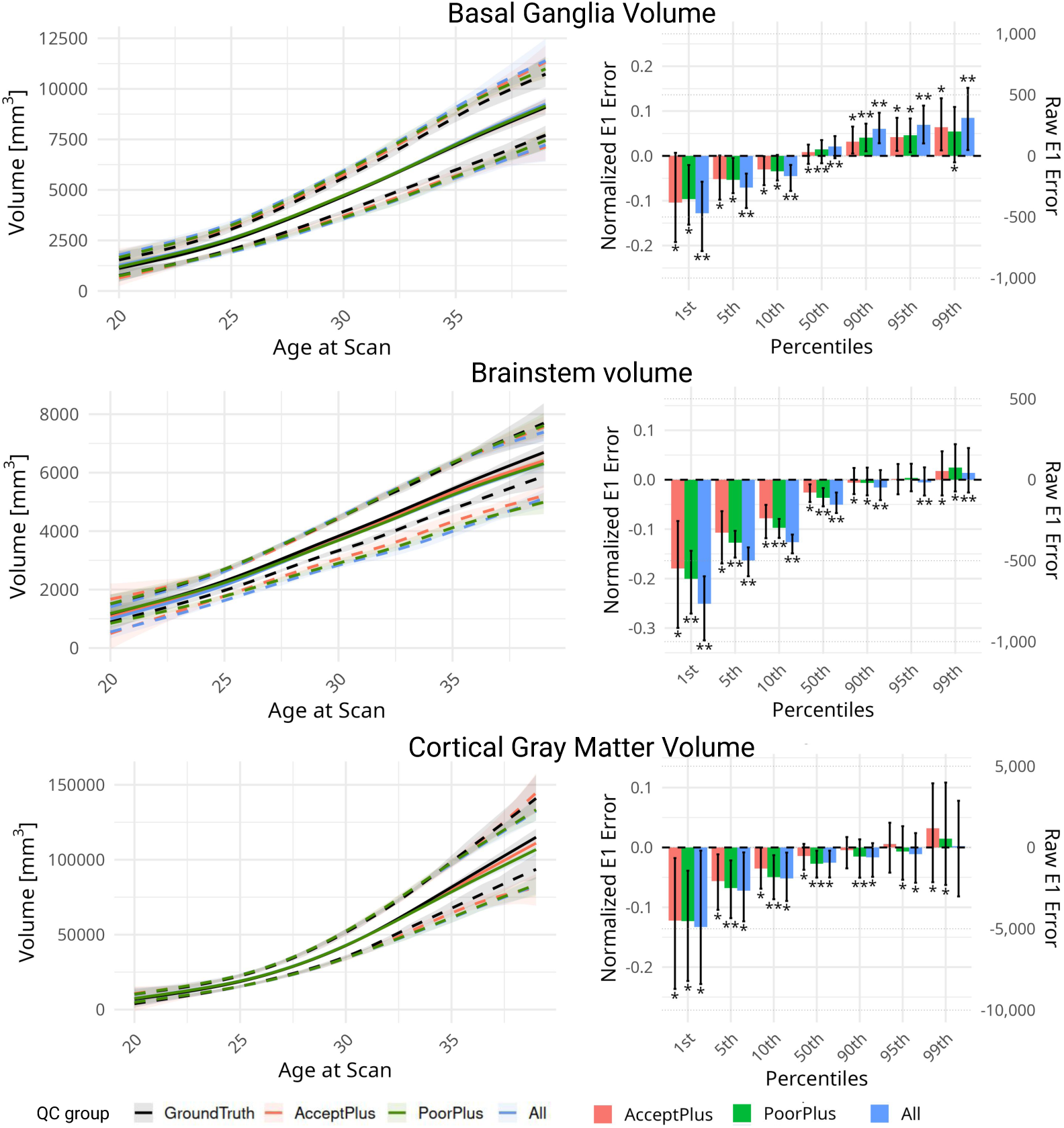
Impact of data quality on normative curves for basal ganglia, brainstem and cortical gray matter volume (from top to bottom). Left column: Bootstrapped curves with 95% confidence intervals for the 5th, 50th and 95th centiles. Right column: E_1_ error distribution with 95% confidence intervals computed for 1st, 5th, 10th, 50th, 90th, 95th and 99th centiles.

The observations the structures here generally follow three kind of trends: the impact of QC is similar to white matter for cortical gray matter, basal ganglia and thalamus volumes — we observe a small but systematic spread; it is similar to lateral ventricles for eCSF — where we see a very large variance across bootstraps, especially in the E1 error —, and similar to cerebellum for the volume of the brainstem — where a strong bias is observed in lower centiles.

### S6 Disentangling the effect of QC and sample size on additional structures

Figure S7 extends the ablation study in Figure 7 and further confirms the results observed. Across all brain structures and sample sizes, there is a bias due to the average QC of the data used to fit the normative model.

Table S2 extends Table 2, also showing p-values, as well as non-significant results.

**Figure S7:**
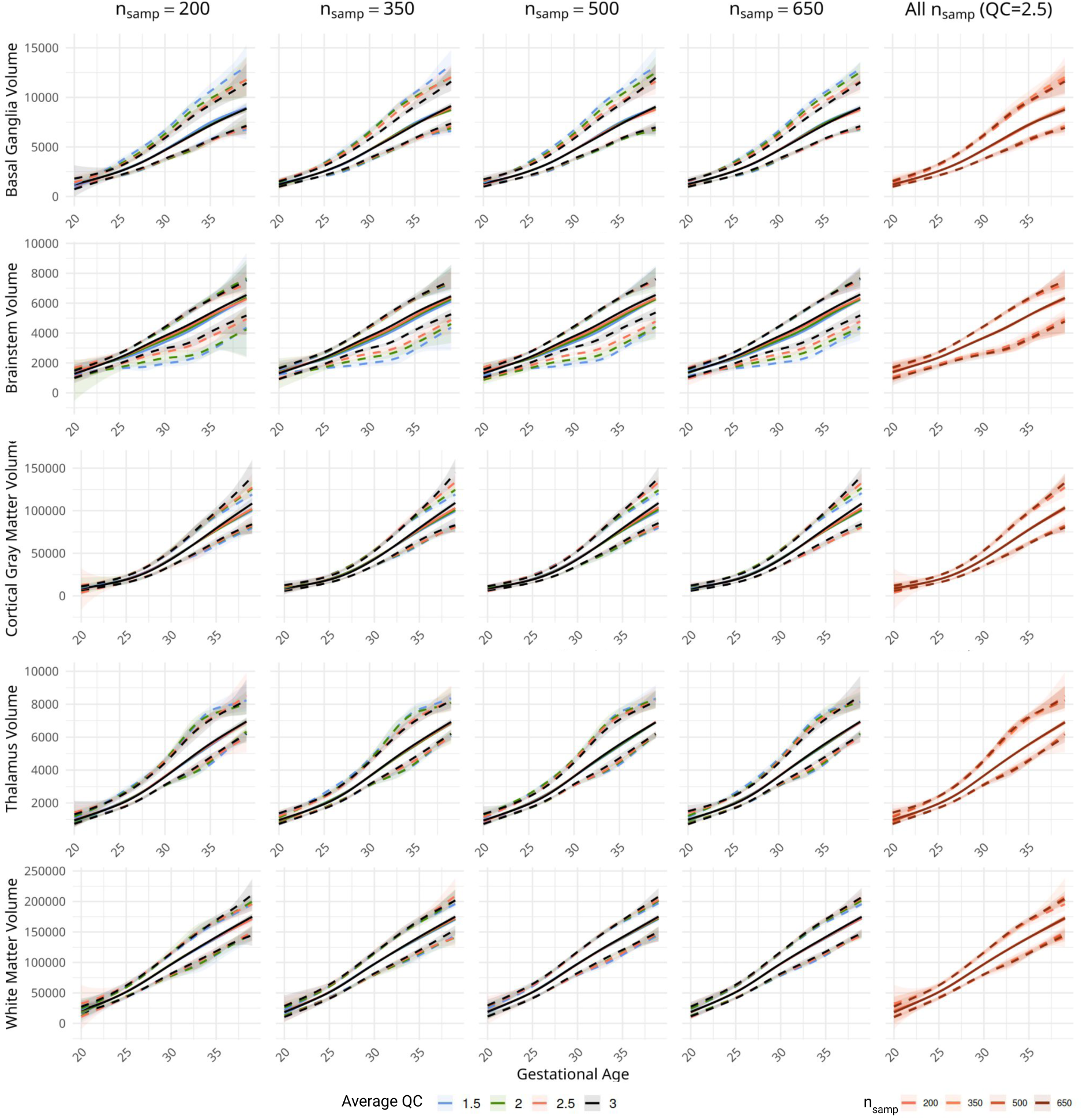
Bootstrapping samples with a given n_samp_ and average quality *q* for additional structures. Extended version of Figure 7.

**Table S2:**
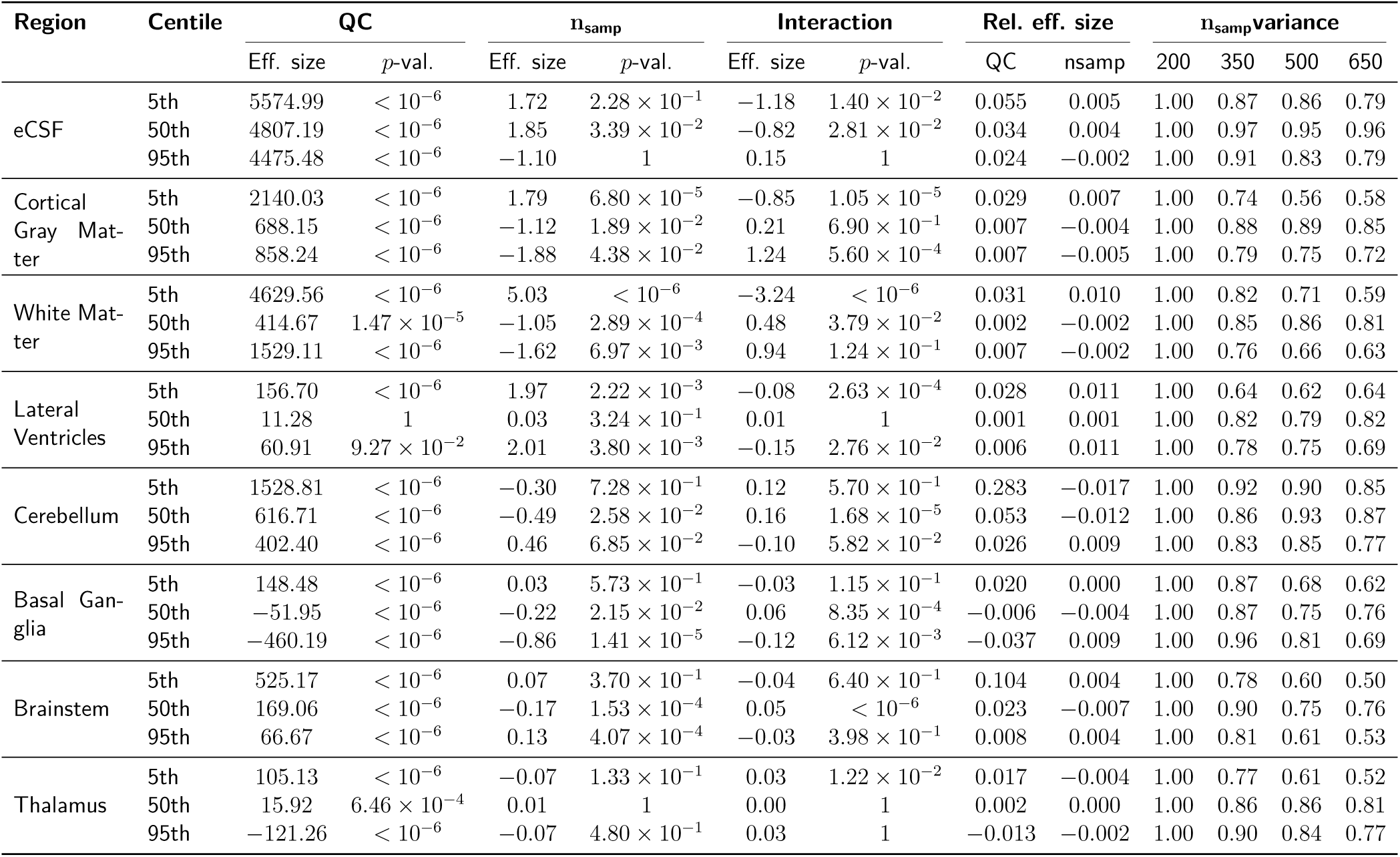
Effect size from a LME model shows that the impact of QC on estimated centiles is consistent, large, and non-uniform. Extended version of Table 2.

### Footnotes

1 https://www.developingconnectome.org

2 https://github.com/fetpype/fetpype

